# Destin: toolkit for single-cell analysis of chromatin accessibility

**DOI:** 10.1101/461905

**Authors:** Eugene Urrutia, Li Chen, Haibo Zhou, Yuchao Jiang

## Abstract

**Summary:** Single-cell assay of transposase-accessible chromatin followed by sequencing (scATAC-seq) is an emerging new technology for the study of gene regulation with single-cell resolution. The data from scATAC-seq are unique sparse, binary, and highly variable even within the same cell type. As such, neither methods developed for bulk ATAC-seq nor single-cell RNA-seq data are appropriate. Here, we present Destin, a bioinformatic and statistical framework for comprehensive scATAC-seq data analysis. Destin performs cell-type clustering via weighted principle component analysis, weighting accessible chromatin regions by existing genomic annotations and publicly available regulomic data sets. The weights and additional tuning parameters are determined via model-based likelihood. We evaluated the performance of Destin using downsampled bulk ATAC-seq data of purified samples and scATAC-seq data from seven diverse experiments. Compared to existing methods, Destin was shown to outperform across all data sets and platforms. For demonstration, we further applied Destin to 2,088 adult mouse forebrain cells and identified cell type-specific association of previously reported schizophrenia GWAS loci.

**Availability:** Destin toolkit is freely available as an R package at https://github.com/urrutiag/destin.

## 1 Introduction

Single-cell assay of transposase-accessible chromatin followed by sequencing (scATAC-seq) is an emerging new technology for the study of gene regulation with single-cell resolution. Unlike conventional regulomics technologies, scATAC-seq measures chromatin accessibility within each individual cell, which circumvents the averaging artifacts associated with traditional bulk population data, yielding new insights into epigenetic regulation at the cellular level. Technologically, Buenrostro *et al.* (2015) adapted the bulk ATAC-seq technology to single cells, utilizing microfluidic device to physically isolate single cells. Cusanovich *et al.* (2015) adopted a two-step “split-and-pool” strategy, where cells undergo several rounds of barcoding procedures to be uniquely labeled. Recently, Preissl *et al.* (2018) developed single-nucleus ATAC-seq (snATAC-seq), adapting the two-step combinatorial indexing strategy to frozen tissues. The Chromium Single Cell ATAC Solution by the 10X Genomics (https://www.10xgenomics.com/solutions/single-cell-atac) can further profile chromatin accessibility across 500 to 10,000 nuclei in parallel. Refer to Supplementary Table 1 for summary statistics of existing platforms and technologies.

The data from scATAC-seq are unique - sparse, binary, noisy with biases and artifacts, and highly variable even within cell types. Supplementary Fig. 1 shows a snapshot of both single-cell and bulk-tissue chromatin accessibility within a 800kb region from chromosome 1, using human monocyte cells and purified bulk samples (Corces *et al.*, 2016), respectively. ScATAC-seq data is sparse and noisy, and the signals share low similarity across cells. On the contrary, bulk signals are highly conserved across the five purified samples. Total depth of coverage in single cells is also greatly reduced, several orders of magnitude lower than bulk. Additionally, since most of the genome has only two copies in a cell and the transposase can cleave and add adaptors only once per copy, for each open chromatin region, at most two sequenceable fragments can be generated and equivalently at most two reads per locus can be obtained after removing PCR duplicates. As such, scATAC-seq data is also highly binary, indicating an open/closed status. Because of the aforementioned uniqueness of scATAC-seq data, neither methods developed for bulk ATAC-seq nor single-cell RNA-seq (scRNA-seq) can be directly applied.

A major advantage of single-cell omics technology is the identification of cellular subpopulations from heterogeneous populations of cells. The structure of a complex tissue is tightly linked with its function, and thus determining the identity and frequency of cell types is crucial and allows for study of disease association at much finer resolution. Several cell-type clustering methods specific to scATAC-seq have been proposed. ScAsAT (Baker *et al.*, 2018) first performs dimension reduction on cell-by-cell Jaccard distance matrix, followed by t-SNE (Maaten and Hinton, 2008) and k-mediods for clustering. scABC (Zamanighomi *et al.*, 2018) begins by clustering cell types using weighted k-mediods, up-weighting cells with higher sequencing depth. To address the sparsity and noise observed in scATAC-seq and to reduce dimension, SCRAT (Ji *et al.*, 2017) and chromVar (Schep *et al.*, 2017) aggregate scATAC-seq read counts across biological features such as transcription factor binding motifs, DNase I hypersensitivity sites (DHSs), genes, or gene sets of interest. This is followed by a further dimension reduction step and clustering.

Here, we propose Destin (*<u>De</u>*tection of cell-type *<u>s</u>*pecific difference in chroma*<u>tin</u>* accessibility), a bioinformatic and statistical framework for comprehensive scATAC-seq data analysis. For cell-type clustering, instead of directly aggregating peaks based on existing genomic annotations, Destin adopts weighted principal component analysis (PCA), with peak-specific weights calculated based on the distances to transcription start sites (TSSs) as well as the relative frequency of chromatin accessibility peaks based on *reference* regulomic data from the ENCODE Project (Consortium *et al.*, 2012). The weights, the hyperparameters, as well as the number of principle components, are cast as tuning parameters and are determined based on the likelihood calculated from a post-clustering multinomial model. The optimal number of clusters is determined using an automated elbow method. Destin is evaluated on scATAC-seq data of 5,800 cells from seven experiments and is benchmarked against existing methods. We show that Destin outperforms the other methods across different data sets and platforms. As a proof of concept, we demonstrate Destin on a scATAC-seq data set of 2,088 adult mouse forebrain cells and identify cell-type specific association of previously reported GWAS loci for schizophrenia.

## 2 Materials and methods

Destin begins with a bioinformatic pipeline to preprocess raw sequencing files and follows with statistical analysis for cell-type clustering and cell type enrichment of previously reported GWAS loci. Specifically, the bioinformatic pipeline includes demultiplexing (for platforms with cellular barcodes), trimming adaptors, mapping reads, filtering blacklist regions, calling peaks in pseudo-bulk samples as aggregates of single cells. After a further quality control procedure, this results in a peak-by-cell chromatin accessibility matrix. Refer to Supplementary Materials for details on bioinformatic analysis. In order to optimally cluster cell types, Destin utilizes both existing genomic annotations and publicly available regulomic data sets as reference to prioritize peaks.

First, we upweight distal regulatory elements (e.g., enhancers) relative to proximal elements (e.g., promoters). Corces *et al.* (2016) showed that distal regulatory elements provide sharper ability for clustering than do proximal elements, and thus, for analysis, focused on accessible chromatin regions 1kb upstream from the TSSs. Similarly, Preissl *et al.* (2018) focused on accessible chromatin regions outside a 2kb window from the TSSs. However, Corces *et al.* (2016) also showed that there is predictive value in the promoter region and meanwhile, Zamanighomi *et al.* (2018) further identified cluster-specific peaks within promoter regions that lead to differential expression. As such, Destin retains all peaks in analysis but opts for a binary weighting scheme: higher weights for distal regulatory elements and lower weights for proximal elements, where peaks are categorized as “distal” or “proximal” based on a 3kb window overlapping the TSSs.

Next, we define a second set of peak-specific weights based on our hypothesis that accessible chromatin regions shared among few cells types are more informative for clustering compared to accessible chromatin regions shared across many cell types. Therefore, we created a reference frequency map for chromatin accessibility peaks using DHS data of 50 to 100 cell lines/types from the ENCODE Project (Consortium *et al.*, 2012), depending on species. The reference DHS frequency was created by calculating the proportion of reference cell types or tissue types containing a DHS peak in each 500bp genomic region. Destin applies a continuous weighting scheme for reference DHS frequency, where higher weights are assigned to accessible chromatin regions with lower reference DHS frequencies.

The two weights for each region/peak are then multiplied to generate a final weight as input for the weighted PCA. Weighted PCA performs dimension reduction in this ultra-high dimensional setting, where binary matrix factorization cannot be run. This is followed by k-means for clustering. The values of weights, as well as the number of PCs, are optimized via grid search to maximize the post-clustering likelihood based on a multinomial model. The number of clusters is determined using the “elbow” method based on the multinomial likelihood. Lastly, a post-clustering step is adopted to re-assign cluster memberships for cells. Refer to Supplementary Materials and Supplementary Algorithm 1 for more details on the proposed method.

## 3 Results

We benchmarked Destin against three existing scATAC-seq methods: scABC, ScAsAT and chromVar. The online GUI by SCRAT takes bam files as input and cannot handle data sets with large number of cells. We began our benchmarks by downsampling bulk ATAC-seq data of purified bulk samples from Corces *et al.* (2016). We varied the number of cell types (2, 4, 6, or 8) and downsampled 50 cells per cell type. Median read depth per cell was also varied, ranging from 3,000 (for combinatorial indexing and 10X Genomics) to 70,000 (for Fluidigm C1). Our results show that performance for all methods increased with number of reads and that Destin outperformed all other methods in almost every scenario, with mean cluster purity shown in Fig. 1A. Next, we benchmarked Destin against the other methods using seven publicly available scATAC-seq data sets (Supplementary Table 2). Destin performed as well as or better than all other methods in terms of cluster purity across all data sets (Fig. 1B), including cluster purity at nearly 100% in five out of seven data sets. For computational efficiency, Destin and chromVar run significantly faster than scABC and ScAsAT across all data sets (Supplementary Fig. 8); for computational capacity, Destin was successfully applied to the 10X Genomics scATAC-seq data of 5k peripheral blood mononuclear cells (see vignettes on GitHub for more details).

**Fig. 1.**
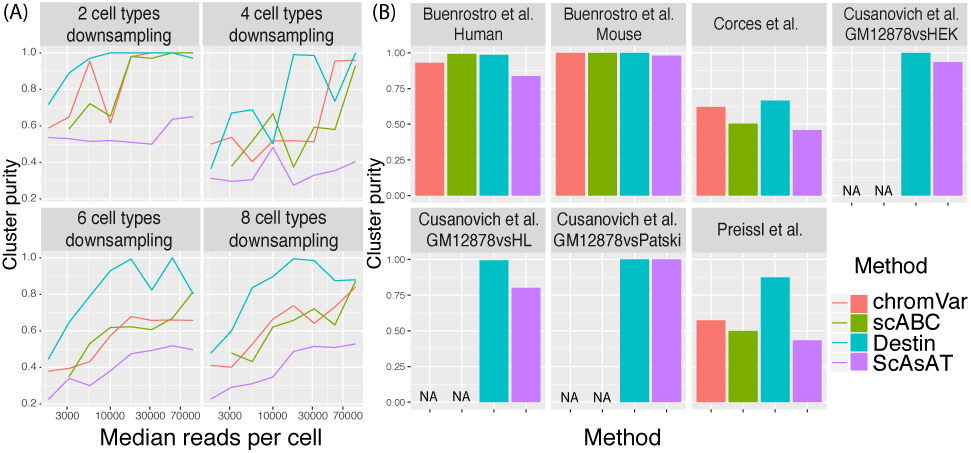
Benchmark results against existing methods via downsampling and empirical data analysis. (A) Purified bulk ATAC-seq data from Corces et al. (2016) were downsampled with different numbers of cell types and different median reads per cell. (B) Cluster results across seven scATAC-seq data sets. chromVAR and scABC cannot be applied to the three data sets from Cusanovich et al. (2015) due to unavailability of required input as bam files.

We further applied Destin to snATAC-seq data of 2,088 cells from adult mouse forebrain (Baker *et al.*, 2018). Destin’s cell-type clustering results, by its default, differed from the results by the original publication – Destin clustered together both the three excitatory neuron subtypes and the related microglia and astrocytes, which we refer to as neuroglia (Supplementary Fig. 9A). Notably, with increased number of cluster, Destin was able to resolve the microglia and astrocytes (Supplementary Fig. 9B).

A key application of single-cell omics technology is to identify specific cell types that are associated with disease. Compared to scRNA-seq, scATAC-seq extends the investigation to non-coding regions including, e.g., promoters and enhancers. In a similar fashion to Skene *et al.* (2018), we determined “cell-type specificity” for each gene, based on the gene annotations of accessible chromatin regions and the clustering results by Destin. The cell-type specificity score was then tested for association with GWAS *p*-values mapped to each gene from three psychiatric studies – schizophrenia (Ripke *et al.*, 2014), major depressive disorder (Wray *et al.*, 2018), and attention deficit hyperactivity disorder (Demontis *et al.*, 2017). We discovered significant association between the schizophrenia GWAS loci and inhibitory neuron 1 by two independent methods MAGMA (de Leeuw *et al.*, 2015) and ECWC (Skene and Grant, 2016) (Table 1). Though an interesting observation whose underlying biological mechanism requires further follow-ups, here by mapping disease risk variants to cell-type-specific regulatory regions, we obtained proof-of-concept identification of possible pathogenic cell types underlying schizophrenia.

**Table 1.** Cell-type enrichment for previously reported GWAS loci. Bonferroni corrected *p*-values are calculated by MAGMA and ECWC for association of mouse forebrain cell types with previously reported GWAS loci for schizophrenia (SCZ), major depressive disorder (MDD), and attention deficit hyperactivity disorder (ADHD).

| Cell Type | MAGMA *p*-value |  |  | ECWC *p*-value |
| --- | --- | --- | --- | --- |
|  | SCZ | MDD | ADHD | SCZ |
| Excitatory neuron | 0.466 | 0.164 | 1 | 0.255 |
| Inhibitory neuron 1 | 0.009* | 0.151 | 0.774 | 0.035* |
| Inhibitory neuron 2 | 0.434 | 0.204 | 1 | 1 |
| Neuroglia | 1 | 1 | 1 | 1 |
| Oligodendrocyte | 1 | 1 | 1 | 1 |

## References

Baker, S. M., et al. (2018). Classifying cells with scasat, a single-cell atac-seq analysis tool. Nucleic Acids Res.

Buenrostro, J. D., et al. (2015). Single-cell chromatin accessibility reveals principles of regulatory variation. Nature, 523(7561), 486.

Consortium, E. P. et al. (2012). An integrated encyclopedia of dna elements in the human genome. Nature, 489(7414), 57.

Corces, M. R., et al. (2016). Lineage-specific and single-cell chromatin accessibility charts human hematopoiesis and leukemia evolution. Nat. genet., 48(10), 1193.

Cusanovich, D. A., et al. (2015). Multiplex single-cell profiling of chromatin accessibility by combinatorial cellular indexing. Science, 348(6237), 910–914.

de Leeuw, C. A., et al. (2015). Magma: generalized gene-set analysis of gwas data. PLoS comput. biol., 11(4), e1004219.

Demontis, D., et al. (2017). Discovery of the first genome-wide significant risk loci for adhd. BioRxiv, page 145581.

Ji, Z., et al. (2017). Single-cell regulome data analysis by scrat. Bioinformatics, 33(18), 2930–2932.

Maaten, L. v. d. et al. (2008). Visualizing data using t-sne. J. mach. learn. res., 9(Nov), 2579–2605.

Preissl, S., et al. (2018). Single-nucleus analysis of accessible chromatin in developing mouse forebrain reveals cell-type-specific transcriptional regulation. Technical report, Nature Publishing Group.

Ripke, S., et al. (2014). Biological insights from 108 schizophrenia-associated genetic loci. Nature, 511(7510), 421.

Schep, A. N., et al. (2017). chromvar: inferring transcription-factor-associated accessibility from single-cell epigenomic data. Nat. methods, 14(10), 975.

Skene, N. G. et al. (2016). Identification of vulnerable cell types in major brain disorders using single cell transcriptomes and expression weighted cell type enrichment. Front. neurosci-switz, 10, 16.

Skene, N. G., et al. (2018). Genetic identification of brain cell types underlying schizophrenia. Nat. genet, page 1.

Wray, N. R., et al. (2018). Genome-wide association analyses identify 44 risk variants and refine the genetic architecture of major depression. Nat. genet., 50(5), 668.

Zamanighomi, M., et al. (2018). Unsupervised clustering and epigenetic classification of single cells. Nat. commun., 9(1), 2410.

