## supplement for "Destin: toolkit for single-cell analysis of chromatin accessibility"

### 1 Bioinformatic pipeline for pre-processing

For publicly available datasets, we downloaded scATACseq fastq files using sraToolkit (Leinonen *et al.*, 2010) following author's provided GEO accession IDs (Supplementary Table 2). Due to the combinatorial indexing technology, Preissl and Cusanovich fastq files were a combination of all cells. Preissl et al. provided decomplexed fastq files, meaning each read pair was identifiable by barcode combination. Cusanovich fastq files, however, were not decomplexed (i.e. reads could not be mapped to specific cells), thus we used their fully processed accessibility matrix, skipping the pipeline altogether.

The first preprocessing step for the Preissl data was to separate the combined fastq files to individual fastq files by cell, using a custom python script. Corces and Buenrostro (Fluidigm C1 technology) fastq files were already separated by cell since the cells were assayed independently. After obtaining fastq file for each cell, both technologies followed the same pipeline.

To align the reads, we first cut Illumina adaptors using cutadapt (Martin, 2011) with –minimum length set to 20. Reads were aligned to respective genome (Supplementary Table 2) using bowtie2 (Langmead and Salzberg, 2012) with setting X2000 to ensure paired reads were within 2kb of one another. Samtools (Li *et al.*, 2009) was used to covert to bam format. Picard tools (Broad Institute, 2018) was then used to perform a series of tasks: SortSam to sort; AddOrReplaceReadGroups to add read groups and index; MarkDuplicates to mark and remove duplicates; and BuildBamIndex to index reads. Samtools was used to remove mitochondrial, unmapped, and chromosome Y reads. Next, due to Tn5 insertion, the starting position of forward reads was adjusted +4, and the end position of reverse reads was adjusted -5. This was performed using a custom awk command. Only aligned reads with map quality over 30 were retained. Finally, aligned reads were sorted by picard SortSam and indexed by samtools.

Peaks were called by MACS2 (Zhang *et al.*, 2008), where all single cells were aggregated to generate a pseudo-bulk sample for peak-calling. We used parameter –nomodel -p 0.01 and thus only peaks with p-value below 0.01 were retained. Peaks were filtered using an ENCODE annotated blacklist file mainly consisting of low-mappability regions and segmental duplication regions (Consortium *et al.*, 2012) : "ENCFF547MET.bed" for mm10 and "wgEncodeDacMapabilityConsensusExcludable.bed" for hg19.

We created the chromatin accessibility matrix (region X cell) by counting the number of reads overlapping each peak for each cell, using countOverlaps from the R package GenomicRanges. Counts were then binarized to create an indicator of whether or not the region corresponding to the peak was accessible. The chromatin accessibility matrix was formatted as a ranged summarized experiment object (R SummarizedExperiment package).

Destin uses the resulting chromatin accessibility matrix as the basis for quality control. Destin retains chromatin accessibility regions where at least 5 unique cells had at least one read overlapping the region. Destin also retain cells with total number of unique accessible chromatin regions within 3 standard deviations (robustly calculated using median absolute deviations) from the median.

The pipeline is publicly available at: `https://github.com/urrutiag/destin`.

### 2 Overview of clustering methods

We compared Destin to 3 other cell type clustering methods specifically designed for scATAC-seq data. To ensure fair benchmarks, we fixed the number of cell types to be the same as the true number. Cusanovich fastq files were not decomplexed (i.e. reads could not be mapped to specific cells). Thus we could not produce bam files which are required by scABC and chromVar.

- scABC clusters cells by weighted k-mediods, where cells with higher coverage have higher weights. To improve performance, scABC then re-clusters cells according to landmarks calculated for each cluster. We compute clustering results using the function scABC from their provided R package scABC with nClusters set to the true cell type number.
- ChromVar computes a bias-corrected deviation for each transcription factor (TF) binding motif, which is followed by Pearson correlation and hierarchical clustering. We found that using the R function hclust followed by cut resulted in poor clustering results. Therefore, we show results where the Pearson correlation is followed by k-means, which produced uniformly better results. We followed the procedure described in Introduction.Rmd from https://github.com/GreenleafLab/chromVAR, using default parameters. Clustering was performed using either R function kmeans or R functions hclust followed by cutree, all from stats package.
- For ScAsAT, we first computed the Jaccard matrix by the proxy R package. K-mediods was computed by R function pam with parameter "diss" set to true and "k" equal to the true number of cell types.
- For Destin, weighted PCA was computed using R function irlba (irlba package). R function kmeans was used for clustering with 100 random starts and "centers" set to the true cell type number.
- The online GUI by SCRAT cannot be applied to the current setting with large number of single-cell bam files as input.

### 3 Weighting proximal v.s. distal regulatory elements

We define proximal regulatory element as within 2kb upstream and 1kb downstream of TSS. Otherwise the accessible chromatin region is defined as distal regulatory element. Destin applies a binary weighting scheme to proximal v.s. distal regulatory elements annotation: A lower weight is assigned to each proximal regulatory element and a higher weight to each distal element:

$$w_j^{TSS} = \begin{cases} w_{proximal}^{TSS}, & \text{if peak } j \text{ is proximal regulatory element,} \\ w_{distal}^{TSS}, & \text{if peak } j \text{ is distal regulatory element.} \end{cases}$$

The two weights ($w_{proximal}^{TSS}$, $w_{distal}^{TSS}$) are cast as tuning parameters and are determined based on the likelihood calculated from a post-clustering multinomial model.

Distance to transcription start site (TSS) is annotated using R function annotatePeakInBatch from the ChIPpeakAnno package. Data set 'TSS.human.GRCh37' is used for model hg19 and 'TSS.mouse.GRCm38' is used for model mm10.

### 4 Weighting common v.s. rare accessible chromatin regions

Destin applies a continuous weighting scheme to common v.s. rare accessible chromatin regions. For each peak $j$ ($1 \leq j \leq m$), a beta distribution maps the corresponding reference DNase I hypersensitive site (DHS) frequency to a nonnegative weight. Specifically, let $DHS_j$ be the percentage of openness across all cell lines and tissues in the reference set (between 0 and 1) for peak $j$, $DHS_j \sim \text{Beta}(\alpha, \beta)$, where $\alpha$ and $\beta$ are hyperparameters shared across all peaks. The probability density function of the beta distribution at $DHS_j$ is used as the weight for peak $j$:

$$w_j^{DHS} = \text{dBeta}(DHS_j, \alpha, \beta).$$

When $\alpha = \beta = 1$, there is no weight; when $\alpha > \beta = 1$, common accessible regions are up-weighted; when $1 = \alpha < \beta$, rare accessible regions are up-weighted. Destin opts to give higher weights to accessible chromatin regions corresponding to rare reference chromatin accessibility frequency. The hyperparameters $(\alpha, \beta)$ of the beta distribution are cast as tuning parameters and are determined based on the likelihood calculated from a post-clustering multinomial model.

To generate the reference chromatin accessibility frequency map, we used DHS data from the ENCODE database (Consortium *et al.*, 2012). For human reference, we used search terms: "bed broadpeaks", "Homo sapiens", "DNase-seq", and "cell line". This resulted in 138 experiments representing 99 cell lines, all from either John Stamatoyannopoulos lab or Gregory Crawford lab. For mouse reference, we used search terms: "bed broadpeaks", "Mus musculus", "DNase-seq", and "tissue". This resulted in 61 experiments representing 27 tissue types. All mouse experiments were from the John Stamatoyannopoulos lab.

The reference DHS frequency was created by calculating the proportion of cell types or tissue types containing a DHS peak in each 500bp genomic region. We created a template representing the entire genome in 500bp bins using R functions tile and GRanges from package GenomicRanges. This was followed by GenomicRanges function overlapsAny (resulting in a binary value) to assign the files to the template. Overlaps were averaged across all files of same type (cell line or tissue), so that no type would be over-represented. Finally, overlaps were averaged across all types.

### 5 Accounting for depth of coverage

Normalization by cell coverage depth prior to principle component analysis would effectively down-weight cells with high depth of coverage, which is the opposite of what (Zamanighomi *et al.*, 2018) have shown to be effective. Other groups (Baker *et al.*, 2017) have found that the first components of dimension reduction are highly correlated with number of accessible chromatin regions per cell. Destin eliminates the correlation between principle components and depth of coverage by applying depth normalization post PCA, which is empirically shown to have better or equal performance in 7 out of 8 data sets (Supplementary Fig. 2). The Corces data set showed improved purity from pre-PCA normalization. Unfortunately, the likelihood was lower, so the likelihood is unable to select pre-PCA normalization

Depth of coverage is calculated by summing the count of accessible chromatin regions by cell from the binarized matrix. Normalization post PCA is performed by dividing each principle component (length nCells) by the depth vector (length nCells).

### 6 Optimization of tuning parameters

To select for the optimal weights and the optimal number of principal components for the weighted PCA, we denote $\boldsymbol{X}$ as the post-QC binarized read count matrix across cells and loci, where $X_{ij}$ indicates the observed binary openness for cell $i$ ($1 \leq i \leq n$) at locus $j$ ($1 \leq j \leq m$). Let $\boldsymbol{C}$ indicate the cluster memberships across all cells, where $C_i = k$ if cell $i$ is from cluster $k$ ($1 \leq k \leq K$). Given the cell type memberships, we reconstruct pseudo-bulk ATAC profile as an aggregate of single cells of the same cell type: $B_{kj} = \sum_{\{i:C_i=k\}} X_{ij}$. Furthermore, we define $P_{kj} = B_{kj} / \sum_{j=1}^{m} B_{kj}$ and this corresponds to the event probabilities in the multinomial distribution:

$$\boldsymbol{X}_i | \boldsymbol{X}, \boldsymbol{C} \overset{iid}{\sim} \text{Multinomial}\left( \sum_{j=1}^{m} X_{ij}, \boldsymbol{P}_{C_i} = (P_{C_i 1}, P_{C_i 2}, ..., P_{C_i m}) \right),$$

where the probability mass function simplifies to

$$p(\boldsymbol{X}_i | \boldsymbol{X}, \boldsymbol{C}) = \left( \sum_{j=1}^{m} X_{ij} \right)! \prod_{\{j:X_{ij} \neq 0\}} P_{C_i j}.$$

Our model assumes that i) the cell-type specific bulk signal is of a pure cell type, i.e., no cell subtypes within, ii) the accessible chromatin regions are independently accessible, and iii) the number of accessible chromatin regions in single cell is much fewer than the number of bulk accessible chromatin regions, leading to binary results. Therefore, we have shown that a multinomial likelihood can be calculated using only unsupervised cluster membership and single cell chromatin accessibility as input. The total likelihood is the product of the individual cell likelihoods, where we assume independence between cells given cell type. This post-clustering likelihood is used to tune parameters and to re-assign cluster memberships across all cells.

We show that the choice of tuning parameters (e.g., number of principal components, promoter/distal weights, reference rare/common weights), which results in higher cluster purity, is data set dependent, and that the tuning parameters can be well approximated using the unsupervised model-based likelihood. After tuning all the parameters, the likelihood can be used to re-assign cluster memberships for individual cells according to maximum posterior likelihood.

Included in the analysis results shown in Supplementary Fig. 3, Supplementary Fig. 4, and Supplementary Fig. 5 were the three scATAC-seq data sets with greater than two cell types. The four data sets with two cell types are not shown, due to saturation of cluster purity. Additionally we show results for the downsampled Corces bulk ATAC-seq data set including all 13 cell types sampled at 50 cells each. Normalized log-likelihood is plotted alongside cluster purity. Normalized log-likelihood is linearly transformed to be on the same scale as cluster purity, for visual clarity.

##### 6.1 Number of principle components

The number of principle components resulting in higher cluster purity was data-set dependent (Supplementary Fig. 3). Cluster purity for the Buenrostro human data set reaches a plateau after 7 principle components, whereas the Corces data set requires 40 principle components to reach a maximum. The Preissl data set and downsampled Corces bulk ATAC-seq data set showed additional diversity. This leads to the conclusion that there is no single choice of number of principle components which is optimal across all data sets.

So the question becomes, how can we decide on the optimal choice in an unsupervised framework, where the cluster purity is unknown since the true cell type identity is unknown. Inspection of Supplementary Fig. 3 shows that our unsupervised metric, the multinomial likelihood, provides an accurate assessment of goodness of fit while not requiring any cell type information. The multinomial likelihood requires only the chromatin accessibility matrix and cell type clustering result as input. We see the multinomial log-likelihood varies in an approximately identical fashion to the cluster purity.

##### 6.2 Peak-specific biological weights

The optimal weighting according to distance from TSS was also data set dependent, as seen in Supplementary Fig. 4. The Corces data set required heavy up-weighting of distal regulatory elements to achieve maximum cluster purity, while the downsampled Corces bulk ATAC-seq data set required less up-weighting. Three of four data sets found improvement by weighting compared to not weighting (weights (1,1)). Similarly, three of four data sets found improvement by including promoter regions compared to excluding promoter regions (weights (0,1)). In general, up-weighting distal regulatory elements (second weight higher) appeared to improve performance. Our unsupervised model-based likelihood follows the rise and fall of cluster purity well. Selection of the weight pertaining to the highest multinomial likelihood resulted in near optimal cluster purity.

Supplementary Fig. 5 shows a similar picture, that optimal cluster purity was achieved at distinct reference chromatin accessibility frequency weights for each individual data set. All data sets showed that up-weighting rare accessible chromatin regions (second weight higher) improved performance while up-weighting common accessible chromatin regions (first weight higher) was detrimental. Three out of four data sets showed improved performance by up-weighting rare accessible chromatin regions compared to not weighting (weights (1,1)). Our metric was able to predict the optimal weight for every data set, achieving maximum cluster purity.

#### 7 Estimating number of clusters

Destin includes five statistical metrics for choosing the number of clusters and by default resorts to the elbow point in the log-likelihood plot. For each method we set the number of clusters search range from 1 to 20. Supplementary Fig. 6 displays the five metrics as they estimate the number of clusters in the Preissl adult mouse forebrain data set. The blue point indicates the chosen number of clusters.

- Silhouette (Rousseeuw, 1987) simultaneously minimizes the within cluster distance while maximizing the between cluster distance. The silhouette metric is defined using terms $a(i)$ representing average distance from cell $i$ to cells in same cluster, and $b(i)$, representing average distance from cell $i$ to cells in the nearest neighboring cluster. Thus defined, silhouette is the ratio $(b(i)-a(i))/\max\{a(i),b(i)\}$, and $k$ is selected as the number of clusters that maximizes average silhouette width across all cells. We implement the method with R function ClusterR::Optimal_Clusters_KMeans, with "criterion" set to "silhouette".

- The gap statistic (Tibshirani *et al.*, 2001) accounts for the tendency to overestimate $n_k$ by using a permutation approach and fitting a null distribution to the WCSSE which naturally decreases as $n_k$ increases. Then the empirical WCSSE is compared to the null distribution with the gap statistic as output. To select the number of clusters the "first SE" method is used which determines the first $n_k$ where the gap statistic is within 1 standard deviation of the following gap statistic. This is implemented using cluster::clusGap followed by maxSE with "method" set to "firstSEmax".

- Alternatively, the distortion method, introduced by Pham *et al.* (2005), accounts for the decrease in WCSSE by comparing to a baseline model incorporating a weight factor based on number of dimensions. We implement the method with R function ClusterR::Optimal_Clusters_KMeans, with "criterion" set to "distortion_fK".

- SCRAT further applies a systematic elbow criteria to the WCSSE (post-aggregation) to estimate $n_k$. They fit WCSSE as a linear spline function of $n_k$ with a single knot. The knot placement which results in the best fit (lowest sum of squared errors) is selected as $\hat{n}_k$.

- Destin uses an unsupervised model-based likelihood to estimate $n_k$. Like WCSSE, the likelihood is non-decreasing with increasing number of clusters. We implement a systematic elbow method similar to that used by SCRAT. but adapted to our model-based likelihood. We fit the likelihood as a linear spline function of $n_k$ with a single knot. The knot placement which results in the best fit (lowest sum of squared errors) is selected as $\hat{n}_k$.

We examined whether our method of determining number of clusters matched with the number of cell types across all seven scATAC-seq datasets. Supplementary Fig. 7 shows that our likelihood-based systematic elbow correctly estimated the number of cell types in six out of seven data sets. The gap statistic correctly estimated the number of cell types in only three out of seven data sets, while the remainder of methods correctly estimated four out of seven data sets. Note that of the eight clusters found by Preissl *et al.* (2018), they annotated eight cell subtypes but five main cell types, which may explain why all methods underestimated in this case.

### 8 GWAS association

A key application of single cell genomics is to identify specific cell types that associated with disease. Skene *et al.* (2018) recently applied scRNA-seq results to schizophrenia GWAS and identified four cell types which were specific for the GWAS association. Their method begins by creation of a cell type specificity matrix. This is essentially the proportion of coverage at the gene level present in each cell type. Specificity across cell types sums to one for each gene. Then the cell type specificity is tested for association with GWAS SNP p-values mapped to each gene. We applied similar methodology utilizing snATAC-seq data from (Preissl *et al.*, 2018), specifically the adult mouse forebrain single cell ATAC-seq experiment.

GWAS studies are typically performed with human subjects. All 3 GWAS sets in our study used hg19. To map mouse accessible regions to human genes, we used R package annotatePeak with setting "select = first" to annotate the nearest gene to the region using data set "TSS.mouse.GRCm38" from the R ChIPpeakAnno package. R package bioMart was used to convert to ensemble geneID to MGI symbol. Next, we mapped mouse to human gene in HGNC annotation using homolog list "HOM_MouseHumanSequence.rpt" from MGI (Smith *et al.*, 2017). Only genes with 1 to 1 homolog mapping were retained. Finally, we used bioMart again to map to Entrez ID, after which accessible regions were aggregated by gene.

After annotation, we aggregate chromatin accessibility counts by gene corresponding to Entrez ID. To create a specificity matrix, we measured the proportion of chromatin accessibility at each gene for each cell type, normalizing so that each gene summed to one across all cell types. We then used MAGMA to test for association with GWAS SNPs. We first called "annotate" with window size 10kb upstream and 1.5kb downstream of transcribed region. Annotated SNPs were then aggregated by gene to a single p-value. The magma model adjusts for covariance in SNP p-values due to linkage disequilibrium. As done by (Skene *et al.*, 2018), we binned the cell type specificity into 40 quantiles, with an additional bin for regions with no accessibility. The cell type specific bin was used as the feature vector for association testing. Finally test for trend between cell type specificity was performed. Magma accounts for gene size, log gene size, gene density, and log gene density. Gene density accounts for the linkage disequilibrium between SNPs in the gene. The model also incorporates correlations between genes.

We validated the results using a second association method Expression Weighted Celltype Enrichment (ECWC) (Skene and Grant, 2016). While magma tests for trend in SNP p-value over increasing cell type specificity, ECWC tests whether the set of significant SNPs are more specific to cell type than are a random set of SNPs. The empirical association is calculated by summing the cell type specificity across all significant SNP genes. The p-value is calculated via permutation, summing the cell type specificity across a random set of genes.

### 9 Computational performance and limits

We benchmarked Destin on the four datasets which provided means to process fastq to bam files: BuenrostroMouse - 192 cells; Corces - 576 cells; Buenrostro Human - 1056 cells; Preissl P56 - 2088 cells (Supplementary Fig. 8). We compared the computational performance across four algorithms: Destin, ChromVar, scABC, and ScAsAT. Computation was performed using an Macbook Pro laptop with a quad-core i7 (utilizing 7 out of 8 threads) with 16 GB of memory. All algorithms began the timing as bam and bed files were read into R, and ended as clustering results were output. To ensure fair benchmarks, we fixed the number of cell type clusters to be the same as the true number. ChromVar was also run in parallel. ChromVar and Destin are the fastest, followed by scABC, and then ScAsAT.

Destin utilizes the excellent Matrix and IRLBA packages in order to increase computational performance. Both packages utilize sparse matrix classes which is entirely appropriate for chromatin accessibility data. In addition to computational speed, memory usage needs to be considered. Destin reads from bam and translates each one to a sparse vector, then aggregates.

In order to further test the capabilities of Destin, we applied to a pair of scATAC-seq data sets from 10X Genomics: peripheral blood mononuclear cells (PBMCs) from a healthy donor, containing 5K and 10K cells respectively. Time to cluster the 5K dataset (5335 cells) was 13 minutes, and the 10K dataset (8728 cells) was 21 minutes. In detail, Destin began by reading the filtered peak bc matrix files, and then proceeded through the clustering pipeline as previously described, including estimation of the number of clusters.

As Destin is an R application, it runs optimally with total $\leq$ 8Gb object size in memory (R uses bits by convention). Thus, Destin has a limit of number of cells it can process in a single batch. Size of the 5K cell / 100K region dataset in R memory was 400 Mb. Destin was able to accommodate 15 replications of the 5K data (75K cells, 5.8Gb) but reached memory error when tested at 20 replications (100K cells, 7.7Gb).

Cusanovich et al. (Cusanovich *et al.*, 2018) recently performed a series of scATAC-seq experiments, encompassing 100K cells across 13 tissue types. The cerebellum set with 5K bp windows contains 600K regions and 2.5K cells, and uses 270 Mb R memory. Destin is not capable of combining all 13 experiments and clustering simultaneously. However, Destin can accommodate the largest of their single experiments.

### 10 Synthetic scATAC-seq data via downsampling

We created the synthetic scATAC-seq dataset from Corces bulk ATAC-seq data. Because the bulk samples were purified using FACS, we assume that they contain reads derived from a single cell type. To create 50 synthesized single cell, we sampled with replacement from among the corresponding bulk cell type bam files. Then we downsampled to a range of 3,000 to 70,000 reads, corresponding to combinatorial barcoding and Fluidigm C1, respectively.

In detail, bulk bam files were created from fastq files, using the same bioinformatics process as described for scATAC-seq. The bam files were subsampled using command "samtools view -b -s", The subsampling rate was set to desired reads / total bulk reads. Desired reads was set to $10^{(\text{nReadsFactor}+\epsilon)}$ where nReadsFactor was varied from log10(3,000) to log10(70,000) and $\epsilon$ randomly sampled from normal(0, 0.5) distribution. All downstream processing for synthetic scATAC-seq followed the same bioinformatic process as described for scATAC-seq.

**Supplementary Algorithm 1:** Algorithm for cell-type clustering by Destin.

**Input :** $\boldsymbol{X} \in \mathbb{R}^{n \times m}$ as post-QC binary chromatin accessibility matrix

$\boldsymbol{N} = \{\sum_j X_{1j}, ..., \sum_j X_{nj}\} \in \mathbb{R}^n$ as read depth across cells

$\boldsymbol{TSS} \in \mathbb{R}^m$ as set of distance to TSS for all loci

$\boldsymbol{DHS} \in \mathbb{R}^m$ as set of frequency of DHS peaks for all loci

**Output:** $\boldsymbol{C}^{opt} \in \mathbb{R}^n$ as vector optimal cluster membership

**for** $l \leftarrow 1$ **to** $n_l$ **do**

- $\boldsymbol{W}_l = f_l(\boldsymbol{TSS}, \boldsymbol{DHS}) \in \mathbb{R}^m$ using the $l$th combination of weights
- $\boldsymbol{X}_w = \boldsymbol{X} \cdot (\mathbf{1}_n \times \boldsymbol{W}_l^T)$
- Singular value decomposition of $\boldsymbol{X}_w$ as $\boldsymbol{UDV}^T$
- $\boldsymbol{X}_{PC} = \boldsymbol{X}_w \boldsymbol{V}$
- $\hat{\boldsymbol{X}}_{PC} = \boldsymbol{X}_{PC} / (\boldsymbol{N} \times \mathbf{1}_m^T)$
- **for** $n_{pc} \leftarrow 1$ **to** $n$ **do**
  - $\hat{\boldsymbol{X}}_{n_{pc}}$ = top $n_{pc}$ projections of $\hat{\boldsymbol{X}}_{PC}$
  - $\boldsymbol{C}_{l,n_{pc}}$ = cell type clustering result of k-means on $\hat{\boldsymbol{X}}_{n_{pc}}$
  - Calculate multinomial likelihood $L(l, n_{pc}) = \sum_{i=1}^{n} \log P(\boldsymbol{X}_i | \boldsymbol{X}, \boldsymbol{C}_{l,n_{pc}})$

$\boldsymbol{C}^{opt} = \boldsymbol{C}_{l^*, n_{pc}^*}$, where $\{l^*, n_{pc}^*\} = \text{argmax} L(l, n_{pc})$.

**Supplementary Table 1:** Summary statistics across different single-cell/nucleus ATAC-seq platforms. Mean number of loci detected per cell, mean number of cells detected per locus, total number of cells, as well as percentages of 0, 1, and $\geq 2$ read counts from the read count matrix were shown.

| Reference | Platform | No. loci per cell | No. cells per locus | Total no. cells | 0 count | 1 count | $\geq$ 2 counts |
|---|---|---|---|---|---|---|---|
| Buenrostro *et al.* (2015) | Fluidigm C1 | 3,521 | 37 | 1,056 | 96.5% | 2.8% | 0.7% |
| Cusanovich *et al.* (2015) | Combinatorial indexing | 1,273 | 4 | 497 | 99.2% | 0.2% | 0.6% |
| Preissl *et al.* (2018) | Single-nucleus ATAC-seq | 1,276 | 28 | 2,088 | 99.1% | 0.9% | 0.0% |

**Supplementary Table 2:** Single-cell and bulk-tissue ATAC-seq data sets adopted. Data sets across different platforms, species, and cell types were collected. Single-cell and bulk-tissue (after downsampling) data were used for benchmark against other existing methods.

| Reference | Platform | Species | Cells | GEO | No. cell types | No. cells | No. peaks |
|---|---|---|---|---|---|---|---|
| Buenrostro *et al.* (2015) | Fluidigm C1 | Human | H1, K562, GM12878, TF-1, HL-60, BJ | GSE65360 | 6 | 1,056 | 184,270 |
| Buenrostro *et al.* (2015) | Fluidigm C1 | Mouse | ES, EML cells | GSE65360 | 2 | 192 | 146,080 |
| Cusanovich *et al.* (2015) | Combinatorial indexing | Combined | GM12878, Patski | GSE67446 | 2 | 497 | 157,770 |
| Cusanovich *et al.* (2015) | Combinatorial indexing | Human | GM12878, HEK293T | GSE67446 | 2 | 714 | 104,260 |
| Cusanovich *et al.* (2015) | Combinatorial indexing | Human | GM12878, HL-60 | GSE67446 | 2 | 656 | 105,233 |
| Corces *et al.* (2016) | Fluidigm C1 | Human | Leukemic cells | GSE74310 | 4 | 576 | 130,448 |
| Corces *et al.* (2016) | FACS + bulk ATAC-seq | Human | Purified hematopoietic cells | GSE74912 | 13 | - | 590,650 |
| Preissl *et al.* (2018) | Single-nucleus ATAC-seq | Mouse | Adult forebrain | GSE100033 | 8 | 2,088 | 132,506 |

**Supplementary Fig. 1:** Snapshot of single-cell and bulk-tissue ATAC profiles. scATAC-seq data from 5 human monocyte cells, aggregate of 5 and 96 single cells, as well as purified bulk samples, are shown. scATAC-seq data are binary, while bulk ATAC-seq data are on an integer scale (max 60 to 100). Sum of scATAC-seq data recapitulates the purified bulk ATAC profile of the same cell type.

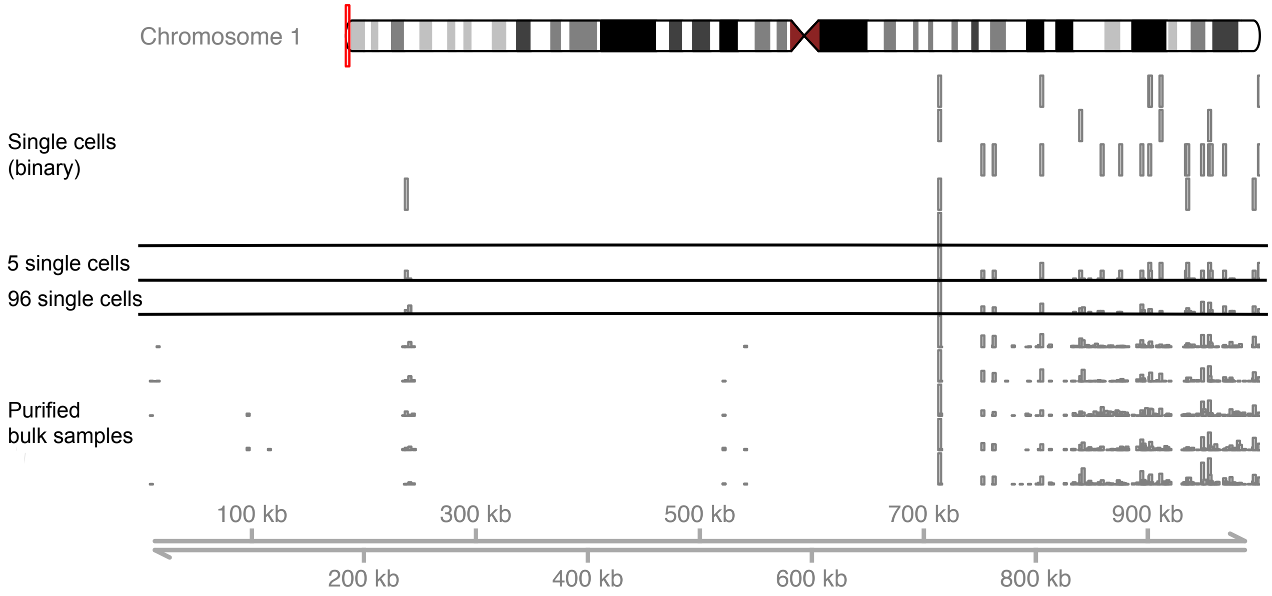

**Supplementary Fig. 2:** Destin eliminates the correlation between principle components and depth of coverage by applying depth normalization post PCA (postPCA). (A) Cluster purity for Destin (postPCA) is compared to a) no adjustment (none) or b) depth normalization prior to PCA (prePCA). (B) First two principle components are not correlated with depth when using post-PCA depth normalization. Mouse data with two cell types from Buenrostro *et al.* (2015) are shown for illustration.

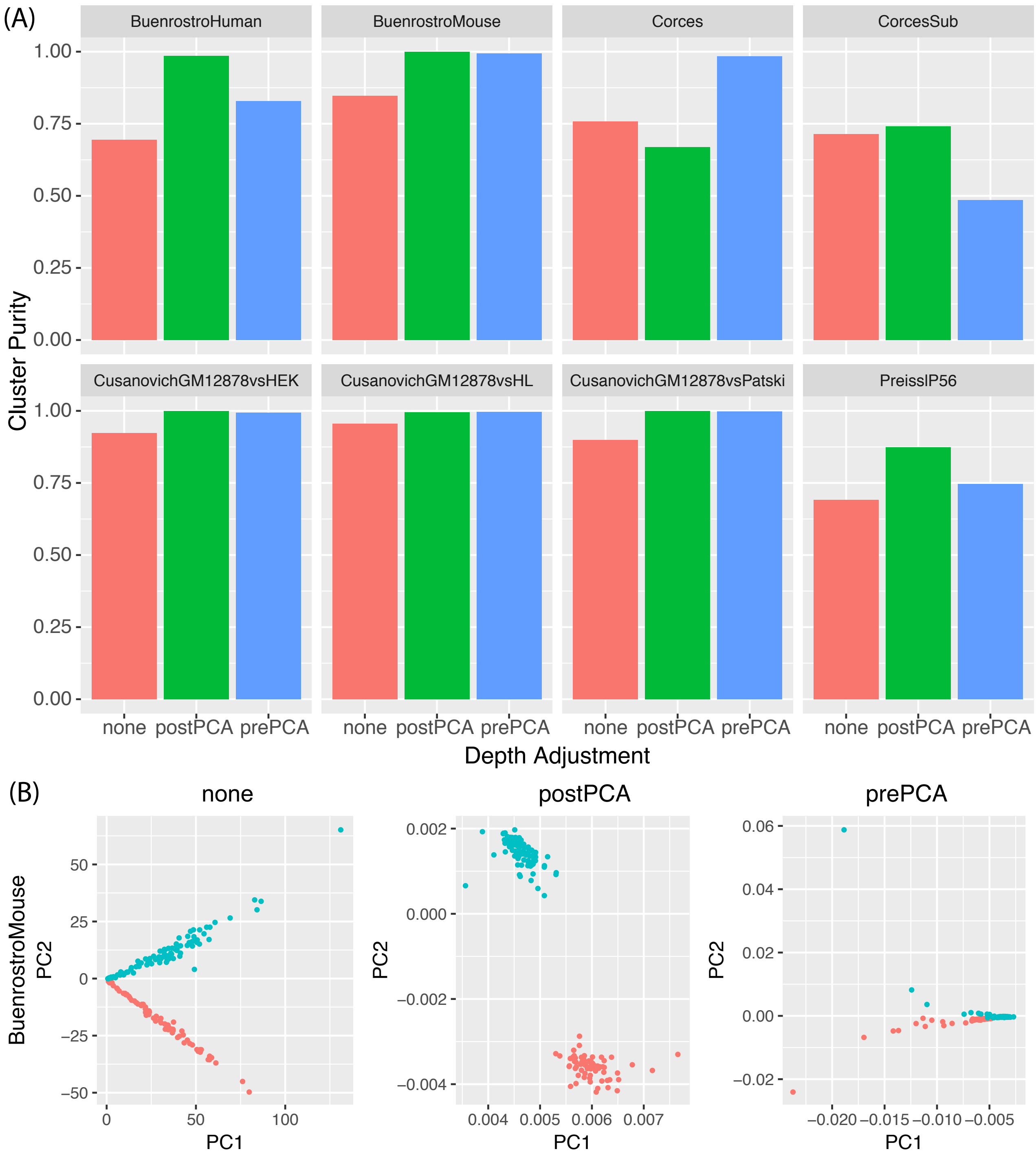

**Supplementary Fig. 3:** Cluster purity and normalized multinomial log-likelihood by number of principle components. Each data point represents a cell type clustering result with no chromatin accessibility region weighting.

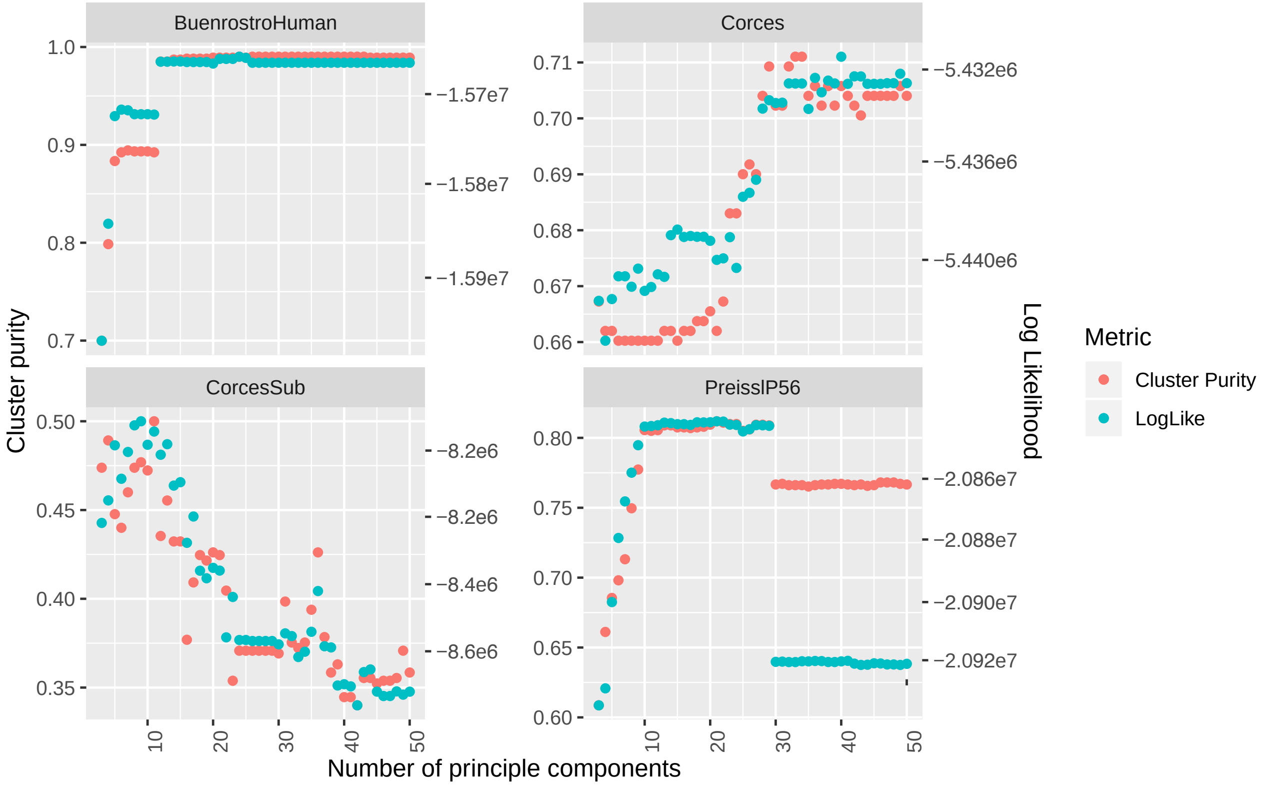

**Supplementary Fig. 4:** Cluster purity and normalized multinomial log-likelihood by weighting according to proximal v.s. distal regulatory element. Each data point represents a cell-type clustering result with 10 principle components and no DHS frequency weighting. Weighting is binary with the first weight corresponding to proximal element and second to distal element. Weights (1,1) correspond to no weighting, while weights (0,1) correspond to removal of promoter regions from the data, and weights (1,0) correspond to removal of distal regulatory elements from the data.

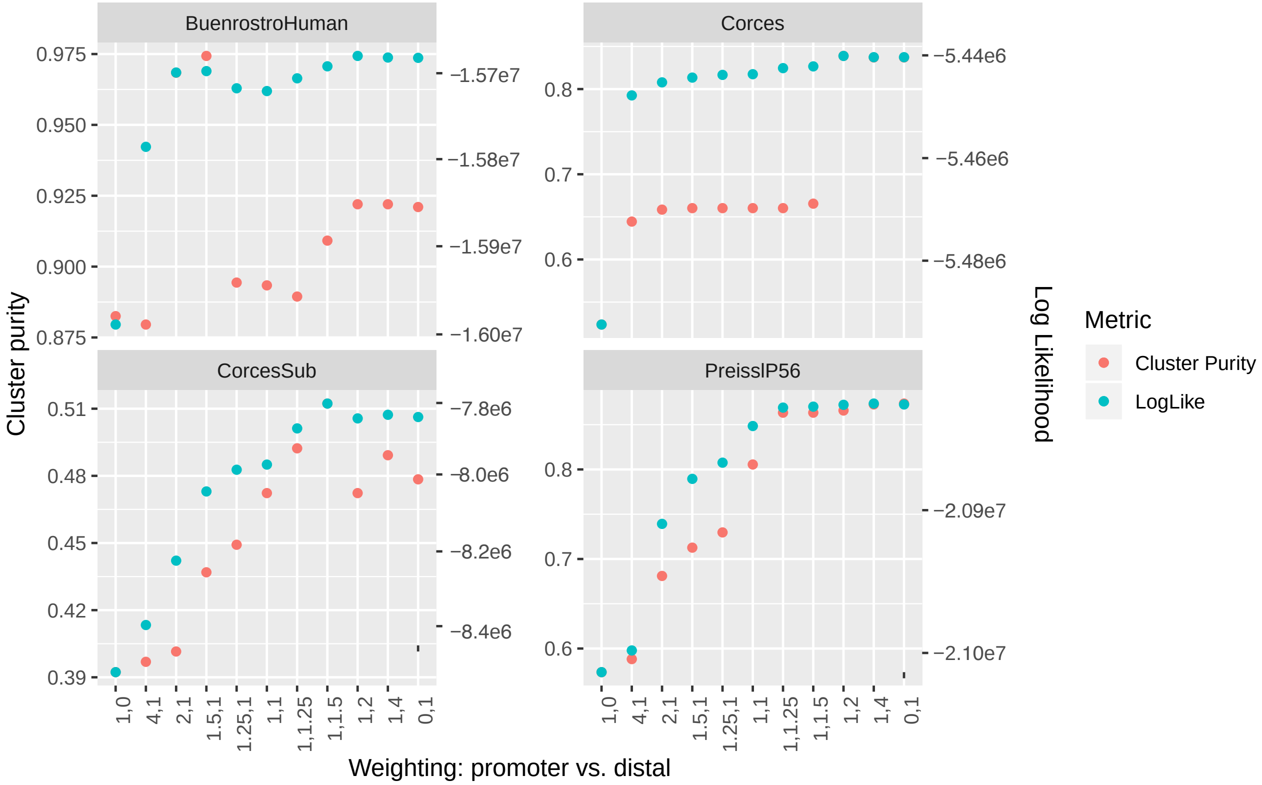

**Supplementary Fig. 5:** Cluster purity and normalized multinomial log-likelihood by weighting according to reference DHS frequency. Each data point represents a cell type clustering result with 10 principle components and no TSS distance weighting. When $\alpha > \beta = 1$, common accessible chromatin regions are up-weighted; when $\beta > \alpha = 1$, rare accessible chromatin regions are up-weighted; Beta(1,1) corresponds to no weighting (uniform weights).

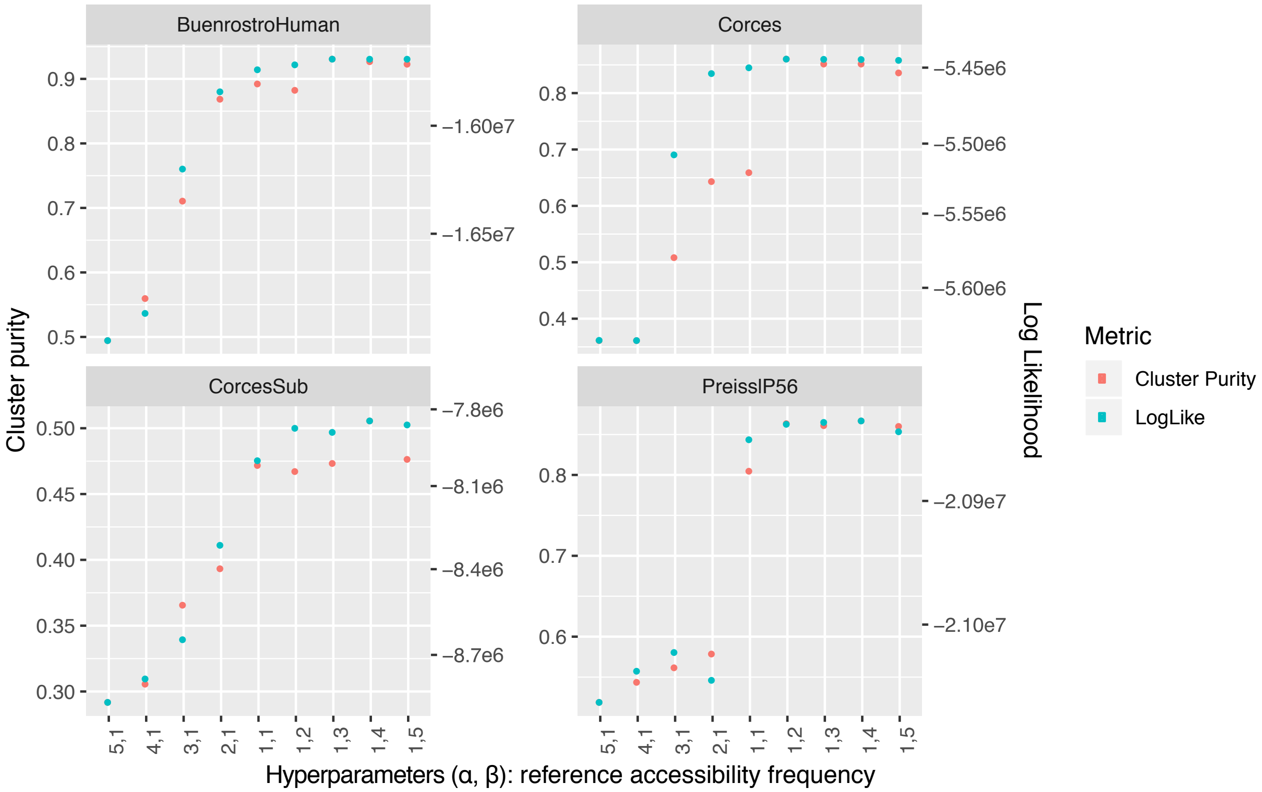

**Supplementary Fig. 6:** Five metrics to estimate the number of unique clusters (cell types) in the Preissl adult mouse forebrain single cell ATAC-seq experiment. Blue point indicates the chosen number of cluster by the given metric. Like WCSSE, the likelihood is non-decreasing with increasing number of clusters. We fit the multinomial log-likelihood as a linear spline function of the number of clusters with a single knot. The knot placement that results in the best fit (lowest sum of squared errors) is selected as number of clusters.

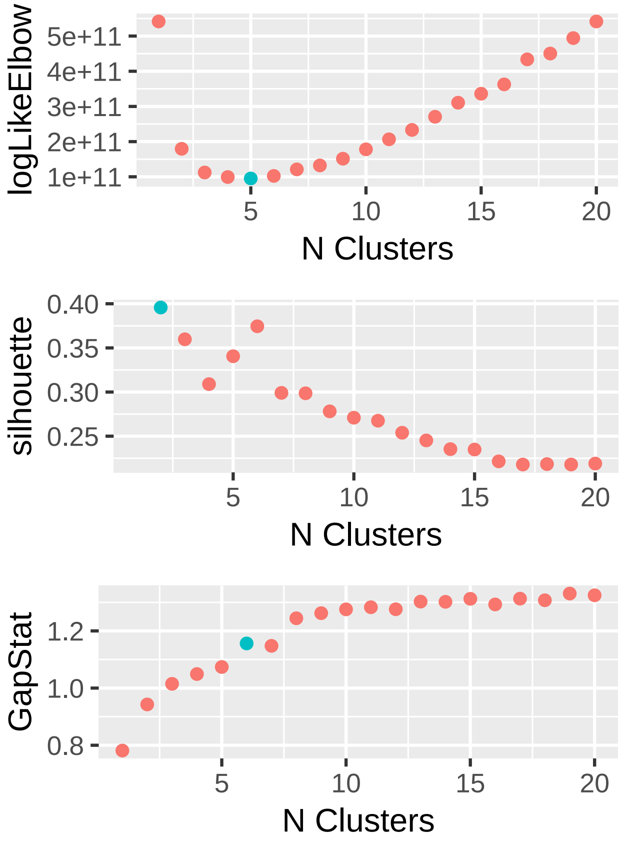

**Supplementary Fig. 7:** Number of clusters estimated across datasets by various methods. Black horizontal line number of cell types provided by original authors

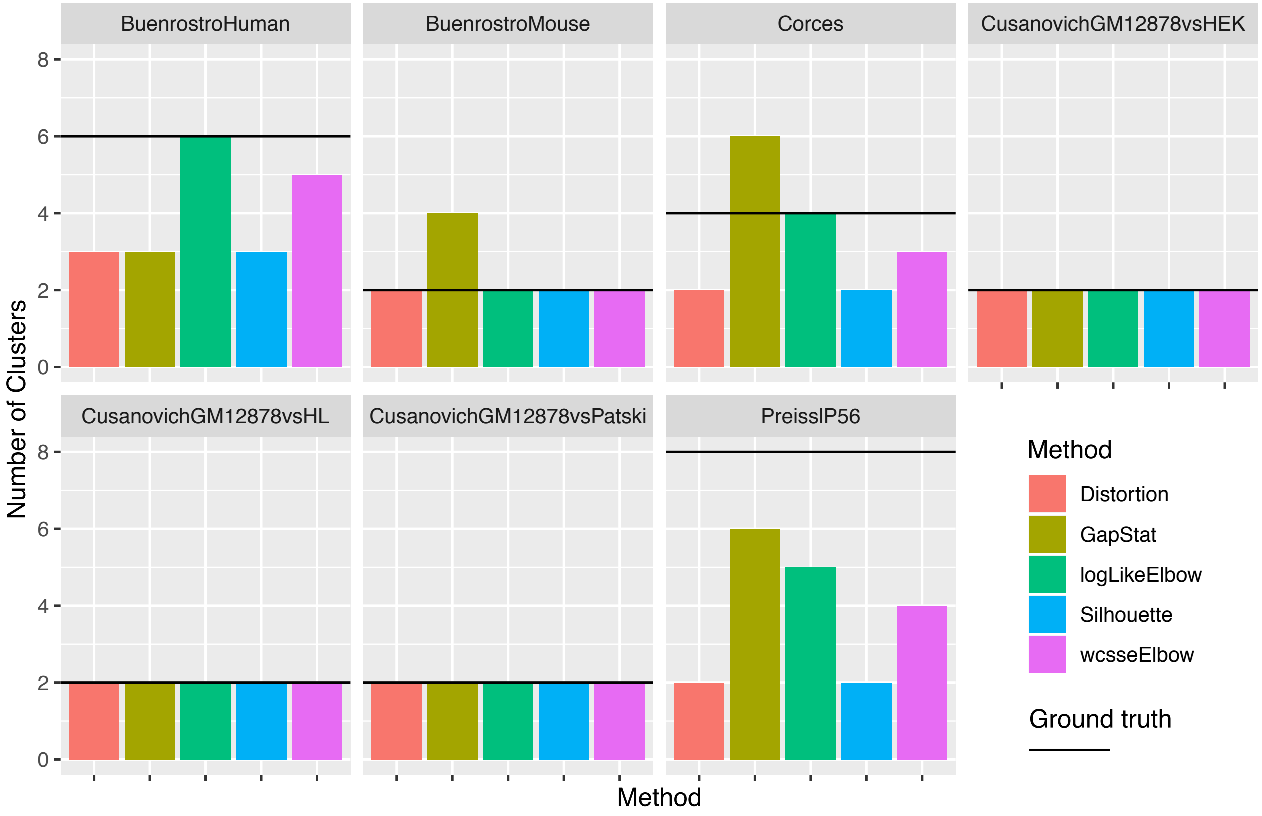

**Supplementary Fig. 8:** Computational benchmarks of Destin and three other algorithms. Four datasets: BuenrostroMouse - 192 cells; Corces - 576 cells; Buenrostro Human - 1056 cells; Preissl P56 - 2088 cells. Number of cells varies by algorithm due to differing QC procedures. Benchmarks performed on Macbook Pro laptop using 7 cores in parallel.

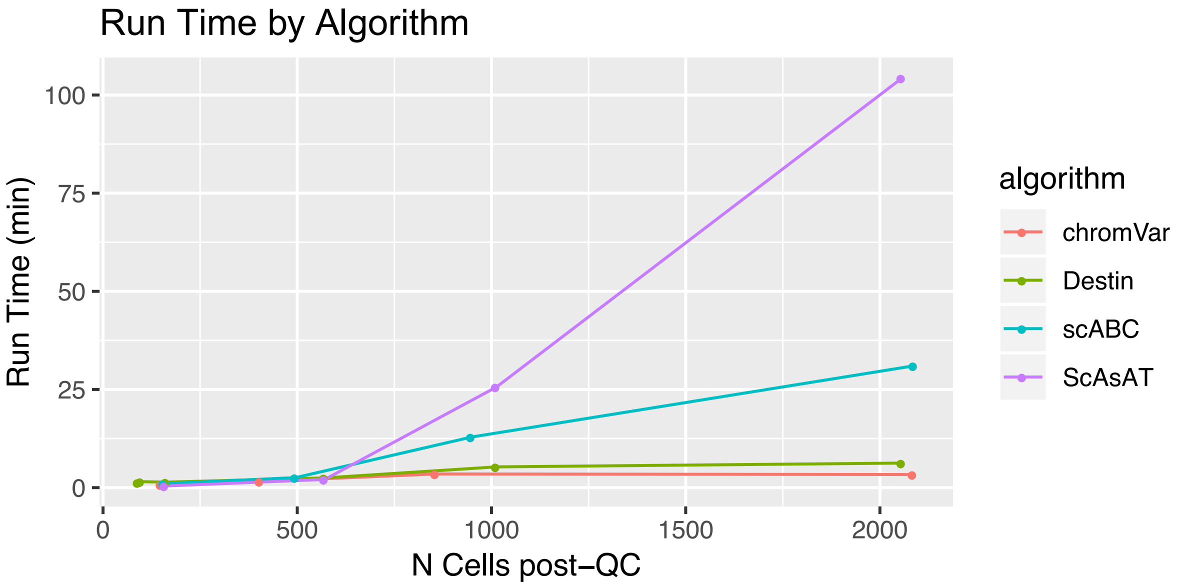

**Supplementary Fig. 9:** t-SNE plot representing the cell type clusters in the Preissl adult mouse forebrain single cell ATAC-seq experiment. For visualization purposes only, as t-SNE is not used by Destin to cluster cells. (A) Destin estimated 5 clusters. (B) Increasing the number of clusters to 6 instead of 5, destin was able to resolve the microglia and astrocyte cell types.

(A)

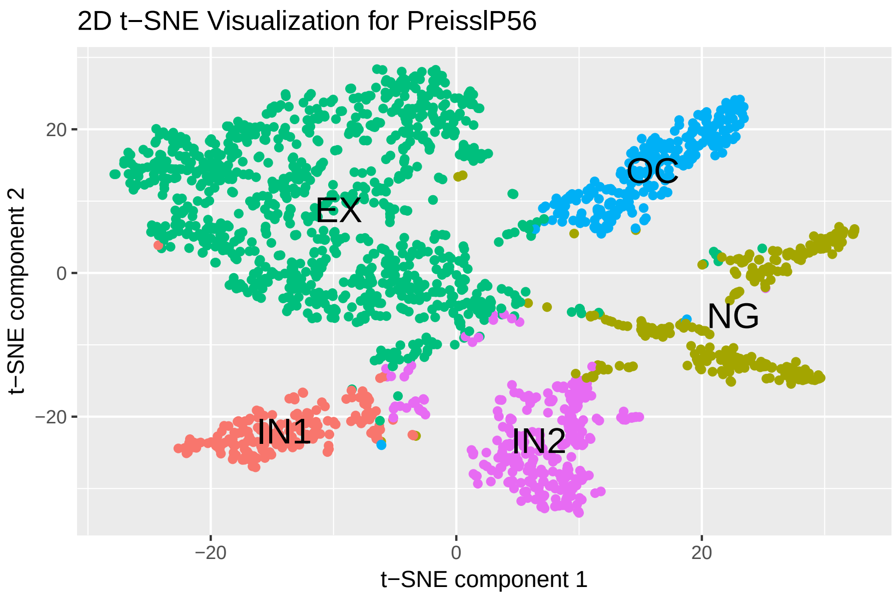


(B)

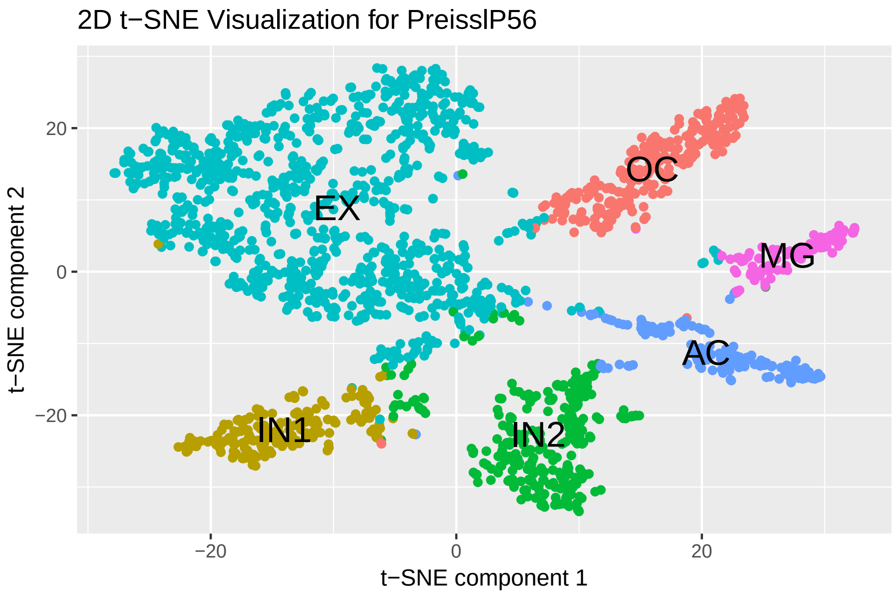

### References


Baker, S. M., *et al.* (2017). Classifying cells with scasat-a tool to analyse single-cell atac-seq. *bioRxiv*, page 227397.

Broad Institute (2018). Picard tools. Accessed: 2018-MM-DD; version X.Y.Z.

Buenrostro, J. D., *et al.* (2015). Single-cell chromatin accessibility reveals principles of regulatory variation. *Nature*, **523**(7561), 486.

Consortium, E. P. *et al.* (2012). An integrated encyclopedia of dna elements in the human genome. *Nature*, **489**(7414), 57.

Corces, M. R., *et al.* (2016). Lineage-specific and single-cell chromatin accessibility charts human hematopoiesis and leukemia evolution. *Nat. genet.*, **48**(10), 1193.

Cusanovich, D. A., *et al.* (2015). Multiplex single-cell profiling of chromatin accessibility by combinatorial cellular indexing. *Science*, **348**(6237), 910–914.

Cusanovich, D. A., *et al.* (2018). A single-cell atlas of in vivo mammalian chromatin accessibility. *Cell*, **174**(5), 1309–1324.

Langmead, B. *et al.* (2012). Fast gapped-read alignment with bowtie 2. *Nature methods*, **9**(4), 357.

Leinonen, R., *et al.* (2010). The sequence read archive. *Nucleic acids research*, **39**(suppl_1), D19–D21.

Li, H., *et al.* (2009). The sequence alignment/map format and samtools. *Bioinformatics*, **25**(16), 2078–2079.

Martin, M. (2011). Cutadapt removes adapter sequences from high-throughput sequencing reads. *EMBnet. journal*, **17**(1), pp–10.

Pham, D. T., *et al.* (2005). Selection of k in k-means clustering. *P. I. Mech. Eng. C-J. Mec.*, **219**(1), 103–119.

Preissl, S., *et al.* (2018). Single-nucleus analysis of accessible chromatin in developing mouse forebrain reveals cell-type-specific transcriptional regulation. Technical report, Nature Publishing Group.

Rousseeuw, P. J. (1987). Silhouettes: a graphical aid to the interpretation and validation of cluster analysis. *J. Comput. Appl. Math.*, **20**, 53–65.

Skene, N. G. *et al.* (2016). Identification of vulnerable cell types in major brain disorders using single cell transcriptomes and expression weighted cell type enrichment. *Front. neurosci-switz*, **10**, 16.

Skene, N. G., *et al.* (2018). Genetic identification of brain cell types underlying schizophrenia. *Nat. genet.*, page 1.

Smith, C. L., *et al.* (2017). Mouse genome database (mgd)-2018: knowledgebase for the laboratory mouse. *nucleic acids res.*, **46**(D1), D836–D842.

Tibshirani, R., *et al.* (2001). Estimating the number of clusters in a data set via the gap statistic. *J. Roy. Stat. Soc. B Met.*, **63**(2), 411–423.

Zamanighomi, M., *et al.* (2018). Unsupervised clustering and epigenetic classification of single cells. *Nat. commun.*, **9**(1), 2410.

Zhang, Y., *et al.* (2008). Model-based analysis of chip-seq (macs). *Genome biology*, **9**(9), R137.
